# Single-objective lattice light sheet microscopy with microfluidics for single-molecule super-resolution imaging of mammalian cells

**DOI:** 10.1101/2025.07.18.665606

**Authors:** Siyang Cheng, Nahima Saliba, Gabriella Gagliano, Prakash Joshi, Anna-Karin Gustavsson

## Abstract

Single-molecule localization microscopy (SMLM) has redefined optical imaging by enabling imaging beyond the diffraction limit, allowing nanoscale investigation into cellular architecture and molecular dynamics. Light sheet illumination enhances SMLM through optical sectioning of the sample, which drastically improves the signal-to-background ratio and reduces photobleaching and photodamage. Lattice light sheet (LLS) microscopy, in which a 2D optical lattice is implemented for light sheet illumination, can provide exceptional sectioning and extended imaging depth when imaging in scattering samples. However, its conventional dual-objective design poses challenges for certain applications. Here, we present an imaging platform which implements LLS illumination with a reflective single-objective geometry (soLLS) inside a microfluidic chip, enabling the use of a single high numerical aperture objective for both illumination and detection, mitigating constraints of a dual-objective setup. We provide quantitative characterization of the propagation properties of the soLLS and demonstrate that it outperforms conventional Gaussian light sheets in terms of useful field of view and sectioning when propagating through scattering samples. Next, by combining soLLS with point spread function engineering, we demonstrate the platform for improved 3D single-molecule super-resolution imaging of multiple targets across multiple cells. The soLLS imaging platform thus expands investigations of nanoscale cellular and intercellular structures and mechanisms into more challenging samples for a wide range of applications in biology and biomedicine.

## Introduction

Fluorescence microscopy is an essential tool for investigating cellular and molecular processes, enabling specific and gentle visualization of subcellular structures^1,2^. Expanding on this capacity, single-molecule localization microscopy (SMLM) has revolutionized the study of biological systems by circumventing the optical diffraction limit, enabling nanoscale examination of cellular architectures and molecular dynamics^3–6^. However, the commonly employed widefield epi-illumination strategy excites the entire sample simultaneously, resulting in a reduced signal-to-background ratio (SBR) due to high fluorescence background from out-of-focus emitters, and increased risk of photobleaching and photodamage to the sample. In SMLM, the reduced SBR deteriorates the localization precision, which in turn degrades the overall resolution of the super-resolved reconstruction. Light sheet (LS) illumination, a strategy where a sheet of light is used to optically section the sample and restrict the illumination to the focal plane, can improve the SMLM performance by increasing the SBR and reducing the photobleaching and photodamage, while still allowing for simultaneous illumination of the entire field-of-view (FOV)^7–10^. Lattice light sheet (LLS) microscopy, which implements a 2D optical lattice for LS illumination, can provide superior penetration depth in thick and scattering samples^11^. In LLS microscopy, the inherent trade-off between optical sectioning and useful FOV of conventional light sheets with a Gaussian beam profile (Gaussian LS) is mitigated by the nature of the optical lattice^11,12^. In combination with SMLM approaches such as DNA points accumulation for imaging in nanoscale topography (DNA-PAINT)^13,14^ and direct stochastic optical reconstruction microscopy (dSTORM)^15^, LLS microscopy has successfully been used for super-resolution imaging of cellular structures^16^. It has also been demonstrated for dynamic imaging of living systems^11^ and for single-particle tracking of plasma membrane molecules^17^.

While widely adopted^11,16,18,19^, the conventional two-objective design of LLS microscopy suffers from several drawbacks. First, the excitation and detection objectives are arranged in an orthogonal geometry, which limits the implementation of high numerical aperture (NA) and short working distance objectives due to physical constraints^11,20^. Second, the objectives are dipped in a media-filled bath, increasing the risk of sample contamination. Third, this design is difficult to combine with solution exchange through microfluidics^21–25^, making DNA-PAINT imaging of multiple cellular structures technically challenging and labor intensive, or requires complex refractive index matching in order to reduce aberrations^14,26–29^. Additionally, the geometry of dual-objective LLS imaging systems can introduce reconstruction artifacts, which require computational correction through post-processing^30^.

In this work, we present a single-objective LLS (soLLS) imaging platform which alleviates drawbacks of the conventional two-objective LLS design. In soLLS, LLS illumination and fluorescence detection are achieved through the same high-NA objective lens, which enables generation of a thin, high-quality, and well-defined LLS while also maintaining high collection efficiency of photons emitted from the sample. By using a reflective micro-optics inside a microfluidic channel to direct the LLS into the sample, the light sheet parameters are decoupled from light sheet steering^31^, in contrast to in e.g. highly inclined and laminated optical sheet (HILO) and its derivatives^32–34^, where the thickness, intensity, position, and depth of the excitation light pattern are coupled and vary depending on the incident angle of the beam. SoLLS thus enables volumetric 3D imaging throughout adherent samples with uniform sectioning. We demonstrate that soLLS outperforms single-objective Gaussian LSs in terms of useful FOV and sectioning when propagating through scattering samples. Furthermore, we demonstrate that soLLS drastically reduces background fluorescence for diffraction-limited cell imaging as well as the localization precision for 2D and 3D single-molecule super-resolution cell imaging compared to conventional epi-illumination. Finally, we demonstrate that soLLS can successfully optically section through multiple cells, enabling improved two-target multi-cell 3D single-molecule super-resolution imaging using sequential DNA-PAINT (Exchange-PAINT)^14^ – enabled and automated through the combination with microfluidics, as well as point spread function (PSF) engineering for nanoscale localization of individual molecules in 3D^35–41^ and deep learning for analysis of overlapping emitters^42^.

Taken together, the improved sectioning capability of the soLLS compared to the conventional Gaussian LS and the easy combination with microfluidics for multi-target super-resolution imaging will enable new biological discoveries by facilitating investigation of more challenging samples.

## Results

### Single-objective lattice light sheet (soLLS) platform design

To break the tradeoff between LS thickness and confocal parameter of a conventional Gaussian LS, a multi-Bessel square LLS^43–46^ with suppressed side lobes was adopted to achieve extended propagation distance while retaining a thin LS for optical sectioning of multiple cells^11,43^. To circumvent limitations of a dual-objective LLS platform, soLLS was designed with a single-objective geometry, allowing LLS formation and fluorescence detection through the same high-NA objective lens (Fig. 1a and Supplementary Fig. 1). The soLLS was generated in a cost-effective and simple manner using a commercially available transmissive binary photomask^47^ for broad implementation (Fig. 1b and Supplementary Fig. 2). A stage-top, microfluidic chip with a metalized nanoprinted insert^31^ was then used both to function as a reflective mirror to redirect the beam for soLLS illumination and to enable precise and automated solution perfusion and exchange using microfluidics. The design of the chip insert micromirror is versatile and can be printed to either reflect the beam horizontally or angled downwards at any angle to facilitate imaging of samples all the way down to the coverslip^31,48,49^. The dimensions of the microfluidic chip can also easily be tuned to accommodate different sample sizes. Additionally, by implementing PSF engineering in the emission pathway, the soLLS setup is capable of extracting 3D information without axial scanning, providing high-precision nanoscale 3D localization of single molecules. In this work, both short- and long axial range double-helix point spread functions (DH-PSFs)^35,36,39^ were used for single-molecule localization and drift correction, respectively (Supplementary Fig. 3).

**Fig 1.**
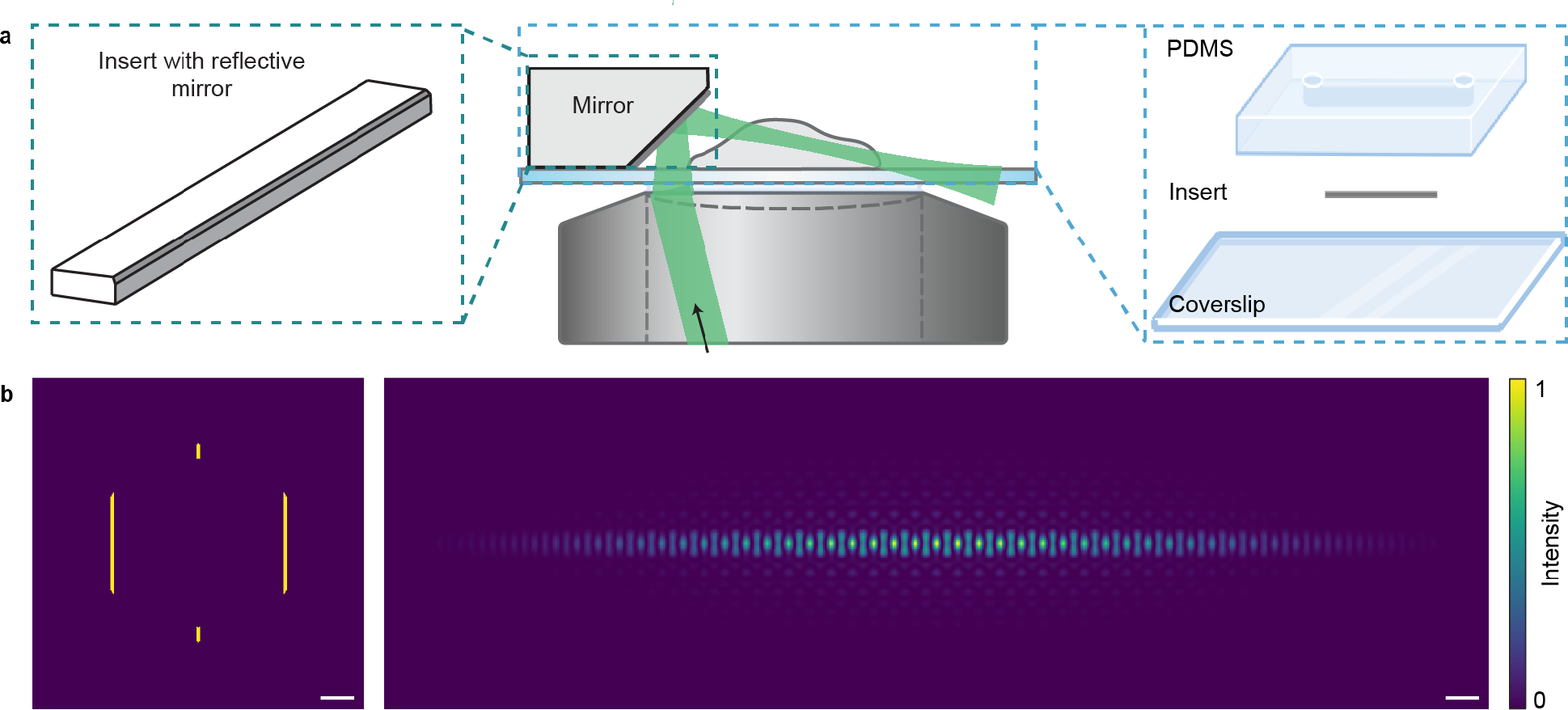
Single-objective lattice light sheet (soLLS) platform design. **a**, The soLLS setup circumvents limitations of the conventional dual-objective LLS approach by implementing lattice light sheet illumination and fluorescence detection through the same high-NA objective lens. A micro-optics incorporated microfluidic chip is used for reflecting and redirecting the soLLS and for automated solution exchange. Schematics not to scale. **b**, Numerical simulations illustrating LLS generation in the setup. The input light field is shaped by a binary transmissive photomask conjugated to the back focal plane (BFP) of the objective, resulting in a LLS formed in planes conjugate to the image plane. Scale bars: left: 1 mm, right: 50 µm.

### Quantification of soLLS characteristics

To quantify the characteristics of the soLLS, its 3D profile was imaged and compared to a single-objective Gaussian LS with comparable thickness at focus (Fig. 2a-c). The soLLS was determined to have a width of

**Fig 2.**
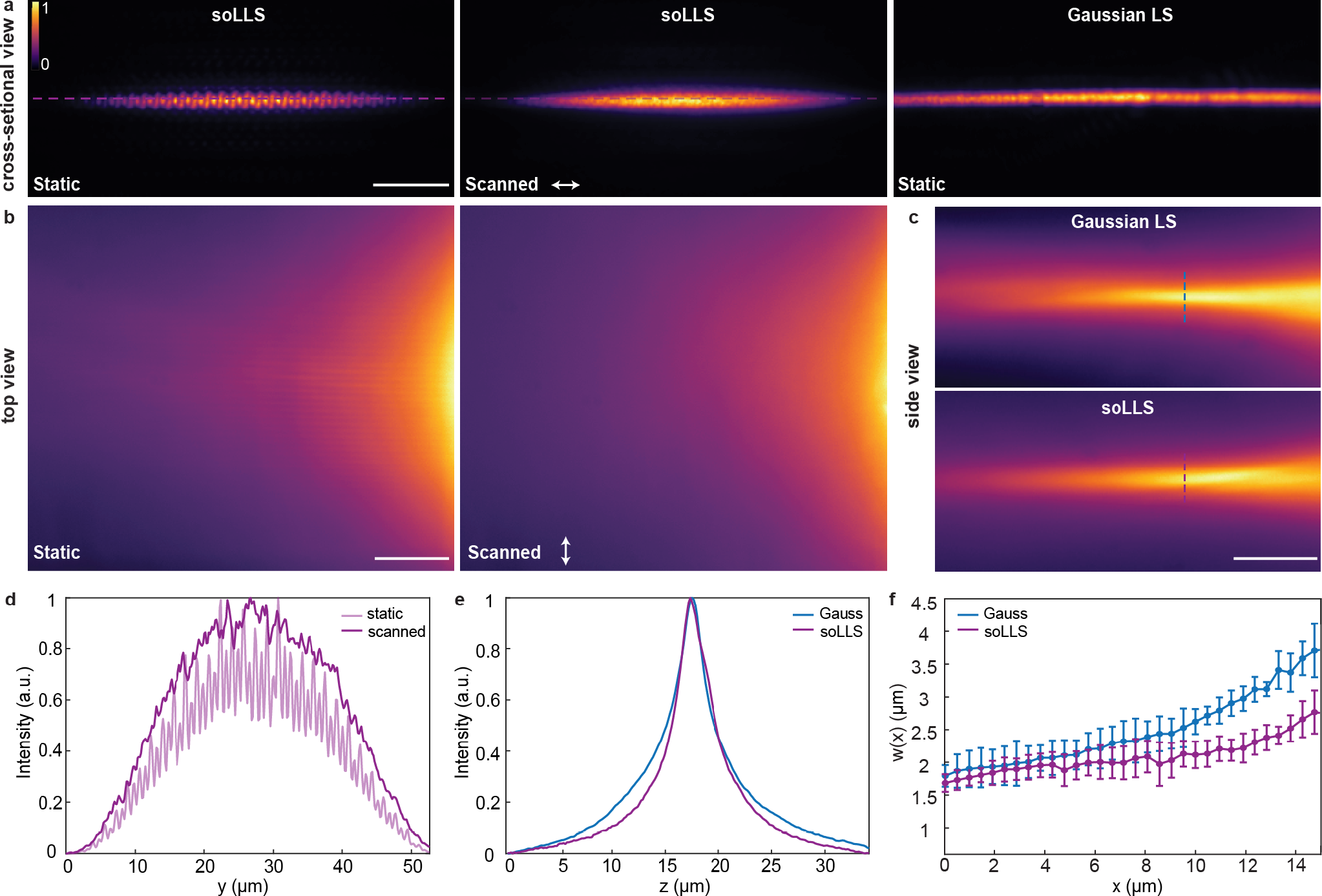
Quantification of soLLS characteristics. **a**, Cross-sectional views of the soLLS profile with and without scanning along the soLLS width direction, and of a Gaussian LS with a similar thickness for comparison. Scale bar: 10 µm. **b**, Top views of the soLLS profile with and without scanning along the soLLS width direction. Scale bar: 10 µm. **c**, Side views of the soLLS and Gaussian LS profiles. Scale bar: 10 µm. **d**, Intensity line scans of the soLLS width acquired from the top views in **b. e**, Intensity line scans of the thicknesses of the soLLS and Gaussian LS measured from the side views in **c. f**, The thicknesses (1/e^2^ radius) of the soLLS and the Gaussian LS as they propagated, measured from the side views in **c**, shown as mean ± standard deviation for n = 3 independent measurements. The colorbar shows normalized intensity.

32.9 µm (1/e^2^ radius) (Fig. 2d). The thickness was determined to be 1.8 µm (1/e^2^ radius) for both the soLLS and Gaussian LS (Fig. 2e), which enabled a direct comparison of their effective ranges (Fig. 2f). Similar to the confocal parameter of Gaussian beams, we here define the range over which the LS thickness remains within a factor of √2 times its beam waist thickness as the effective range of the LSs. The soLLS was determined to have an effective range of 27.9 ± 4.4 µm (mean ± standard deviation, n = 3), thus providing a 1.5-fold improvement over the Gaussian LS that had an effective range of 19.2 ± 0.7 µm (mean ± standard deviation, n = 3) (Fig. 2f and Supplementary Fig. 4). These results demonstrate that the soLLS maintains its thin profile over a larger effective propagation range compared to a conventional Gaussian LS when initialized with a similar beam waist radius, thereby extending the imaging FOV within which desired optical sectioning is achieved.

Unlike in HILO and similar designs^32–34^, where the thickness, intensity, position, and depth of the excitation light pattern are coupled and vary depending on the incident angle of the beam, the soLLS can be repositioned in 3D over the entire imaging FOV in a linear fashion using beam steering units^50^, where the soLLS dimensions are decoupled from the steering (Supplementary Fig. 5). Thus, consistent sectioning performance can be achieved across different regions of interest. Steering axially and in the soLLS plane was implemented using galvanometric mirrors, where 0.01 V applied to the galvanometric mirrors resulted in a translation of the soLLS of 0.64 µm in the axial (z) direction and 0.35 µm in the direction within the soLLS plane (y), respectively. A tunable lens was used to shift the soLLS focus, where 10 mA applied to the tunable lens corresponded to a focus shift of 2.41 µm (Supplementary Fig. 5).

### SoLLS propagates more robustly in scattering environments compared to a Gaussian LS

Self-reconstructing (or self-healing) is another defining characteristic of non-diffracting beams, referring to the ability of such beams to regenerate their initial profiles under free propagation after being disturbed by an obstacle^51^. This property has been experimentally demonstrated for various beam types^52–55^ and it is of particular interest in LS microscopy, as imaging of biological samples inherently suffers from scattering and aberrations due to sample heterogeneity and refractive index variations. LLS, physically described as a coherent superposition of a linear array of Bessel beams^11^, is expected to preserve this feature to an extent depending on beam type and experimental implementation.

To assess the performance of the soLLS propagation in scattering environments, we quantified the ability to localize individual emitters in a scattering environment generated by mixing a solution of fluorescent beads and dye in agarose gel when sectioned with soLLS compared to a Gaussian LS (Fig. 3a,b). Our results demonstrate that, compared to a Gaussian LS, while having comparable numbers of localizations close to the beam focus, illumination with the soLLS yielded a much larger number of localizations at distances further from the beam focus, suggesting that the soLLS is more robust to aberrations caused by scattering environments (Fig. 3c and Supplementary Table 1). In the region furthest from beam focus (Region 5, Fig. 3c), soLLS offers a 370-fold improvement in the number of detected emitters compared to the Gaussian LS (Supplementary Table 1). Furthermore, the SBR values and uncertainties of the localizations obtained with soLLS illumination outperformed those obtained with Gaussian LS illumination in all regions (Fig. 3d,e). Together, these results show that the soLLS behaves more robustly in complex and scattering environments, offering benefits for cell and tissue imaging where imaging at multiple microns of light sheet penetration is required.

**Fig. 3.**
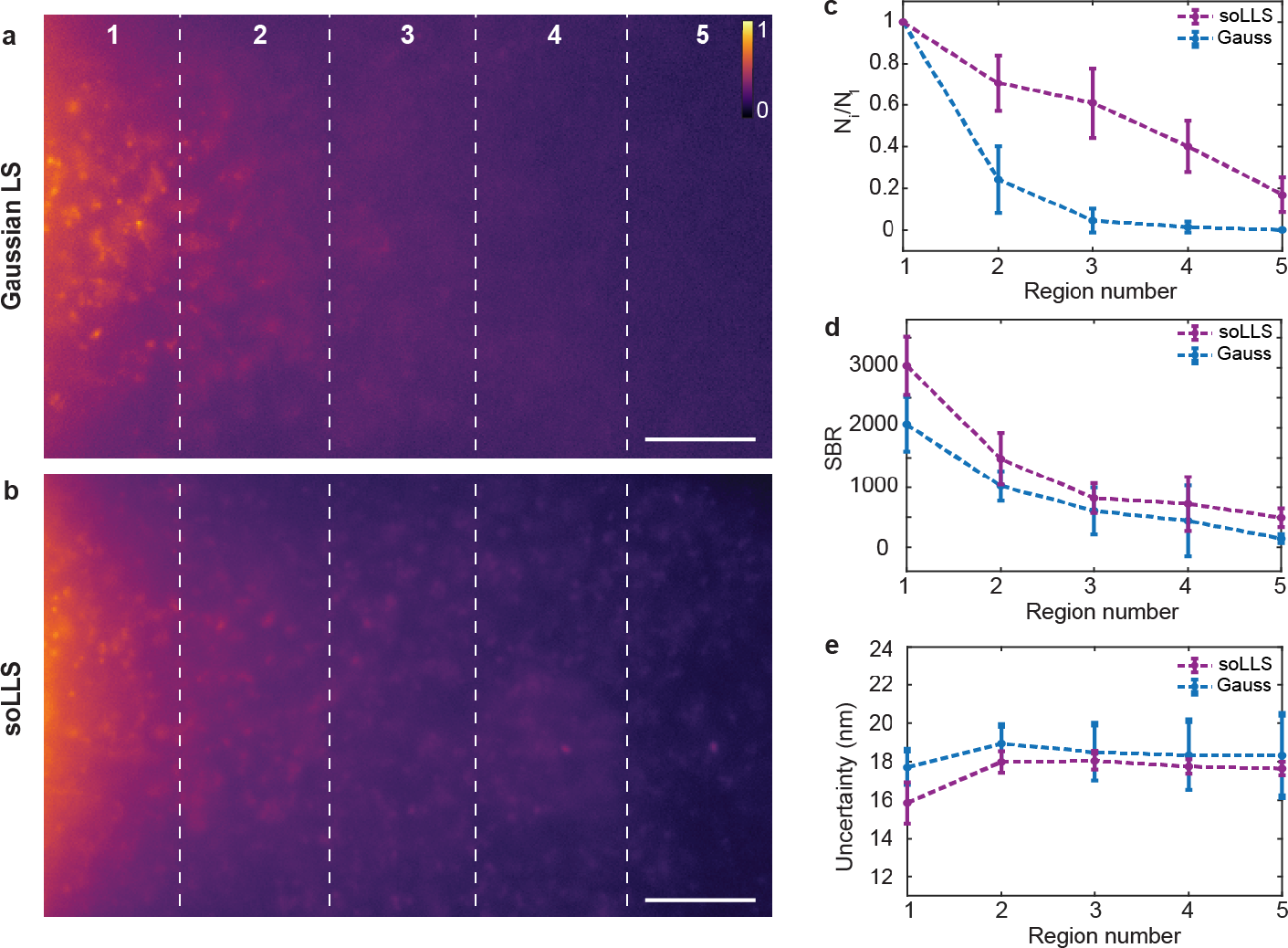
SoLLS propagates more robustly in scattering environments compared to a Gaussian **LS**. Representative images of the **a**, Gaussian LS and **b**, soLLS propagating through scattering environments created by mixing fluorescent beads and dye in an agarose gel. Localizations from the images were binned into 5 rectangular regions of equal area to compare statistics, with region 1 being the closest to the beam focus. Scale bars: 10 µm. The colorbar shows normalized intensity. **c**, Comparison of the number of localizations in each region, N_i_, normalized by the number of localizations in region 1, N_1_. **d**, Comparison of the signal-to-background ratio (SBR) of the localizations in each region. **e**, Comparison of the localization uncertainty in each region. Error bars in **c-e** represent the mean ± standard deviation of the corresponding values across n = 9 fields of view (FOVs). Taken together, soLLS improves both the number of detected particles, the SBR, and the localization precision compared to the Gaussian LS when imaging through scattering samples.

### SoLLS improves the signal-to-background ratio and 3D localization precision for cellular imaging

The quality of single-molecule data is often degraded when imaging throughout thick samples due to out-of-focus background fluorescence. To demonstrate that soLLS effectively optically sections thick samples, such as mammalian cells, and thereby improves the SBR for both diffraction-limited and super-resolution imaging, lamin B1 in the same U2OS cells were imaged back-to-back with soLLS and widefield epi-illumination. This data showed that soLLS improved the SBR by up to 7.5-fold for diffraction-limited imaging (Fig. 4a) and up to 3.9-fold for single-molecule imaging (Fig. 4b). Technical replicates for diffraction-limited imaging (Supplementary Fig. 6, n = 3 cells) and for single-molecule imaging (Supplementary Fig. 7, n = 4 cells) demonstrated reproducible performance of the soLLS across multiple samples. Next, the improvement for 3D single-molecule imaging was quantified by imaging of lamin B1 using a short-range DH-PSF with soLLS and widefield epi-illumination. The data showed that while having more signal photons, with median values of 4320 photons/localization with soLLS illumination compared to 2654 photons/localization with epi-illumination, the median background photons were drastically reduced to 16.1 photons/pixel with soLLS illumination compared to 25.5 photons/pixel with epi-illumination (Fig. 4c). This resulted in improvements in both lateral (xy) and axial (z) localization precisions, with median values of 11.9 nm laterally and 18.1 nm axially with soLLS illumination compared to 18.1 nm laterally and 27.4 nm axially with epi-illumination. Technical replicates demonstrate reproducible improvements of soLLS for 3D single-molecule imaging across multiple samples (Supplementary Fig. 8, n = 4 cells).

**Fig. 4.**
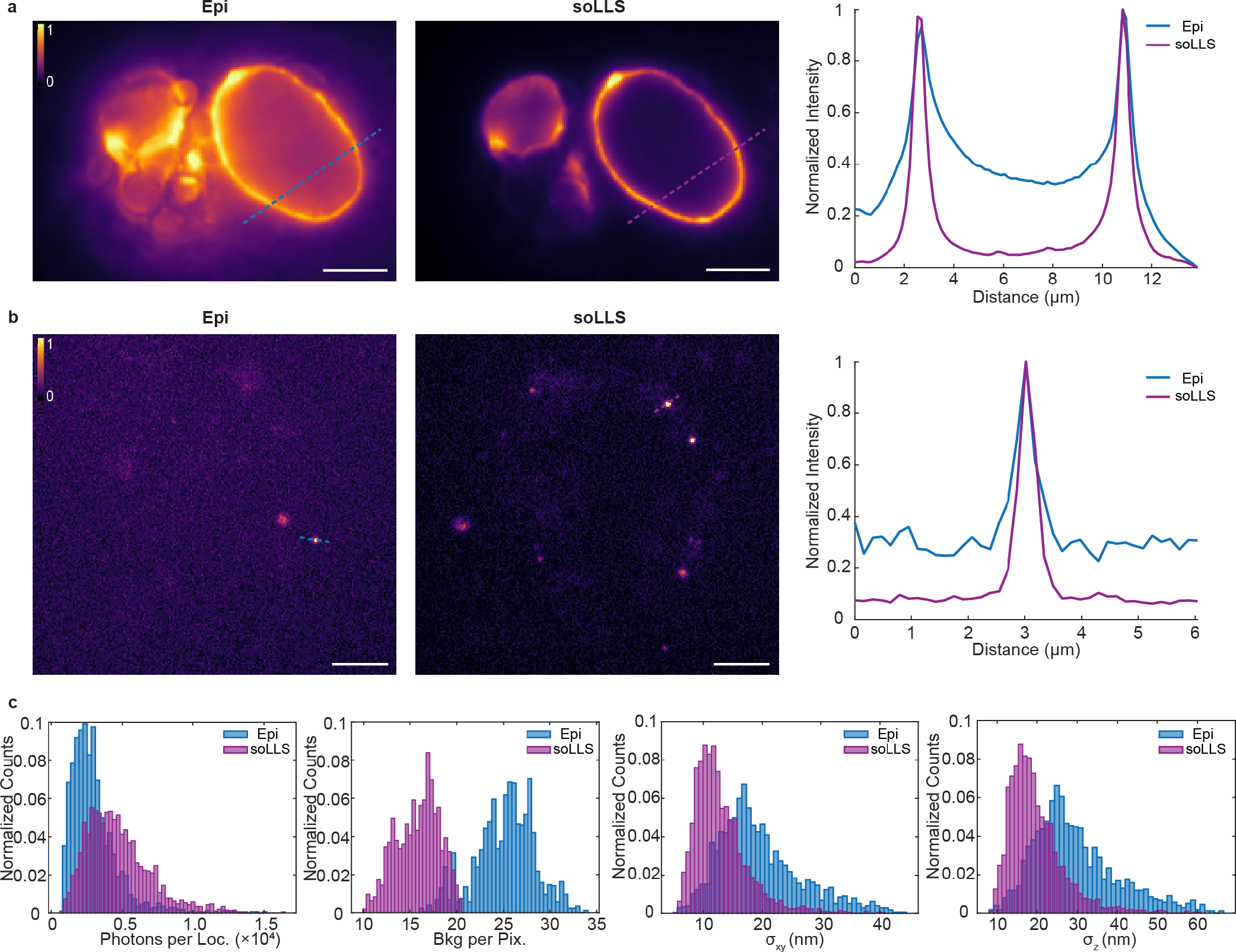
SoLLS improves the signal-to-background ratio (SBR) and 3D localization precision for cellular imaging. **a**, Diffraction-limited images of lamin B1 in U2OS cells acquired with widefield epi-(Epi) or soLLS illumination. Scale bars: 5 µm. The colorbar shows normalized intensity. Graph shows a comparison of the normalized intensity across line scans in the same cell. **b**, 2D single-molecule images of DNA-PAINT labeled lamin B1 in U2OS cells acquired with epi- or soLLS illumination. Scale bars: 5 µm. The colorbar shows normalized intensity. Graph shows the normalized intensity across line scans of single molecules. **c**, Histograms showing comparisons of signal photons per localization, background photons per pixel, lateral (xy) localization precision, and axial (z) localization precision of the localized emitters in the same cell under either epi- or soLLS illumination, demonstrating that soLLS illumination drastically reduces background photons compared to epi-illumination, which results in improved localization precision for 3D single-molecule super-resolution imaging.

### SoLLS provides sectioning for improved 3D single-molecule super-resolution imaging of multiple targets and multiple cells

Light sheet sectioning through multiple cells is typically degraded due to scattering, limiting its benefits for single-molecule imaging of e.g. cell-cell junctions. Furthermore, many questions in biology and biomedicine require imaging and accurate correlation of multiple cellular structures or molecular distributions at the nanoscale, which can be hard to achieve using multi-color imaging or manual solution exchange. With soLLS, multi-target 3D single-molecule super-resolution imaging is made simple by the single-objective configuration in combination with the microfluidic chip for automated solution exchange. Having also shown that soLLS propagates more robustly through complex samples and that it provides excellent optical sectioning of cells, we next set out to demonstrate soLLS for 3D single-molecule super-resolution imaging of several cellular structures and throughout multiple cells.

First, soLLS was demonstrated for two-target Exchange-PAINT imaging of lamina associated protein 2 (LAP2) and mitochondria (TOMM20) using the microfluidic chip for automated solution exchange (Fig. 5a-c). The resulting Fourier ring correlation (FRC) resolutions in the xy/xz/yz planes were found to be 19.2/23.4/23.4 nm for LAP2 and 24.2/30.8/28.5 nm for TOMM20 (Fig. 5d and Supplementary Fig. 9), demonstrating that soLLS achieves multi-targeting imaging with high spatial resolution in 3D (Fig. 5e).

**Fig. 5.**
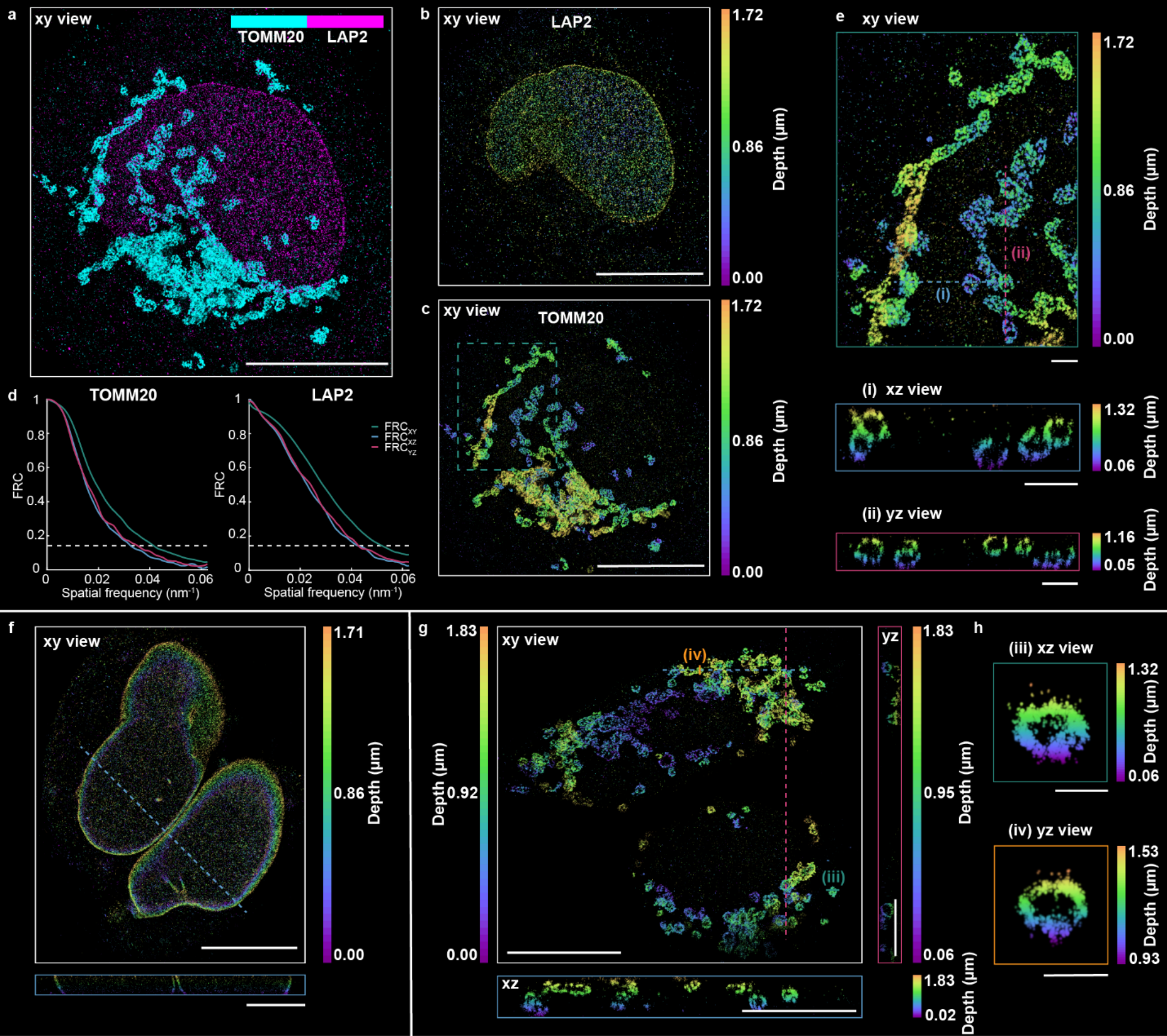
SoLLS provides sectioning for improved 3D single-molecule super-resolution imaging of multiple targets and multiple cells. **a**, Two-target 3D super-resolution reconstruction of LAP2 and mitochondria (TOMM20) in the same U2OS cell, colored by target. Scale bar: 10 µm. Separate renderings of **b**, LAP2 and **c**, TOMM20, colored by depth. Scale bars: 10 µm. **d**, The resulting FRC resolutions in the xy/xz/yz planes for LAP2 and TOMM20. **e**, Zoomed in views of the mitochondria in the dashed box in **c**. The xz and yz views are shown for a 200 nm thick y-slice in the xz plane and a 200 nm thick x-slice in the yz plane at the dashed lines (i) and (ii) in the xy view. Scale bars: 1 µm. **f-g**, SoLLS sectioning and 3D super-resolution imaging in two adjacent cells shown for **f**, lamin A/C and **g**, mitochondria (TOMM20). Scale bars: 10 µm in the xy views and 5 µm in the cross-sectional views. **h**, Zoomed in views of mitochondria cross sections acquired along the solid lines (iii) and (iv) in **g**, 300 nm and 200 nm thick, respectively. Scale bars: 500 nm.

Next, leveraging the enhanced propagation performance of the soLLS, multiple cells were successfully sectioned and imaged across the same FOV, which is typically challenging for conventional Gaussian LSs of similar thickness. Simultaneous 3D single-molecule super-resolution imaging of two cells, labeled for either lamin A/C (Fig. 5f) or mitochondria (Fig. 5g,h), was achieved with resulting FRC resolutions in the xy/xz/yz planes of 33.1/45.1/45.0 nm for lamin A/C and 25.1/33.6/29.5 nm for the mitochondria (Supplementary Fig. 9), demonstrating that the high spatial resolution is retained even when illuminating through multiple cells.

## Discussion

In this work we presented soLLS, an imaging platform which implements a reflected single-objective lattice light sheet generated by a simple printed photomask in combination with a micro-optic incorporated into a microfluidic chip for reflection of the light sheet and for easy and automated solution exchange. The single-objective geometry enables LLS illumination and fluorescence detection through the same high-NA objective, eliminating limitations of the conventional dual-objective LLS design. We provided a quantitative experimental characterization of the soLLS profile in 3D and compared its propagation properties to a Gaussian LS with the same thickness. We demonstrated that soLLS provides optical sectioning over 1.5-fold longer propagation distances compared to a conventional Gaussian LS, due to reduced divergence of its main lobe during propagation. Moreover, we demonstrated that the soLLS propagates more robustly in highly scattering environments, yielding up to 370-fold more localizations further away from the beam focus. Furthermore, we demonstrated an up to 7.5-fold SBR improvement for diffraction-limited imaging and 3.9-fold SBR improvement for single-molecule imaging. The excellent sectioning by soLLS improved the 3D single-molecule localization precision from 20 nm to 13 nm laterally and from 30 nm to 20 nm axially by reducing out-of-focus background photons. Next, we demonstrate soLLS for 3D single-molecule super-resolution imaging of two-targets through Exchange-PAINT with automated solution exchange, which is typically challenging with conventional LLS designs. Finally, we showcase simultaneous 3D single-molecule super-resolution imaging of multiple cells across the FOV, which can typically not be sectioned well with regular Gaussian LSs.

The soLLS imaging platform can be applied to address a large variety of biological and biomedical questions. The extended imaging FOV enabled by the soLLS platform facilitates imaging of e.g. intercellular proteins in multiple cells or mitotic cells, and of molecular 3D distributions in cell-cell junctions^56–59^. The soLLS platform is live-cell compatible and can be used with conventional stage-top incubators, enabling fast, precise, and gentle 3D single-particle tracking in live cells^60–63^. The microfluidic chip simplifies the process of introducing buffers and solutions to the sample, which facilitates multi-target Exchange-PAINT imaging extended beyond the two-target imaging demonstration presented in this work^14,31^. Additionally, it enables well-controlled and reversible introduction of drugs or other stimuli for live-cell studies^64–68^. SoLLS can in the future easily also be implemented with spatial light modulators for flexible and adaptable LLS design^11,69,70^. Taken together, soLLS offers a powerful, flexible, and cost-effective approach for improved 3D single-molecule super-resolution imaging of mammalian cells. SoLLS can be utilized for various single-molecule imaging applications for improved nanoscale investigation of cellular architectures and molecular dynamics.

## Methods

### Single-objective lattice light sheet (soLLS) imaging platform

The soLLS platform was built around an inverted microscope (IX83, Olympus) with a high-NA oil-immersion objective (UPLXAPO100X, 100x, NA 1.45, Olympus) (Supplementary Fig. 1). The excitation laser (560 nm, 1000 mW, MPB Communications) was spectrally filtered (FF01-554/23-25, Semrock) and circularly polarized using a polarizer (LPVISC050MP2, Thorlabs) and a quarter-wave plate (Z-10-A.250-B-556, Tower Optical). The laser was collimated and expanded to a desired spot size using multiple lens telescopes. The LLS was generated with a custom-designed photomask (HTA photomask) conjugated to the back focal plane (BFP) of the objective. The photomask had multiple patterns for flexible tuning of the LLS dimensions and accommodation to different laser spot sizes in the optical path. The characteristics of the LLS are determined by the dimensions of the slits pattern. In this study, we employed a pattern which has a distance of 2.4 mm between the outer slits and slits width of 0.08 mm (Supplementary Fig. 2). Two galvanometric mirrors (GVS011, Thorlabs) and a tunable lens (EL-3-10-VIS-26D-FPC, Optotune) were aligned conjugated to the BFP of the objective lens to function as beam steering units. In addition to positioning the soLLS, the galvanometric mirror used for translating the soLLS in the soLLS plane was scanned with a sine signal at 20 Hz with 0.1 V amplitude to generate a uniform illumination profile. Additionally, a dithering mirror (SP30Y-AG, Thorlabs) conjugated to the image plane was used to dither the soLLS with a sine signal at 100 Hz with 0.1 V amplitude to remove any residual shadowing artifacts^71^. After propagating vertically after the objective, the soLLS was reflected and redirected by a microfluidic chip with an insert that functioned as a reflective mirror. The microfluidic chip was fabricated as detailed in ref^31^. In brief, an insert with custom-designed dimensions and an angled side wall was printed using two-photon polymerization, and the side wall was metalized to function as a reflective mirror. The insert was assembled with a PDMS base and a glass coverslip to serve as the sample chamber.

For comparison between the soLLS and the conventional Gaussian LS, a single-objective Gaussian LS was generated with a cylindrical lens and aligned to have a similar thickness at focus as the soLLS. The Gaussian LS shares the beam steering units with the soLLS, and several flip mirrors were used for switching between the two LS modalities (Supplementary Fig. 1). Additionally, a widefield epi-illumination path was integrated into the imaging platform for comparisons between different illumination modalities and overall flexibility and versatility of the setup.

The fluorescence emission from the sample was collected by the same objective lens and focused by the tube lens inside the microscope. The collected emission light then passed through the first lens of a 4f-system before being split by a dichroic mirror (T660lpxr, Chroma) into two channels. Each of the channels had a second 4f lens that focused the light onto an EMCCD camera (iXon Ultra 897, Andor). The 4f systems provide access to the Fourier plane in each channel to enable phase modulations for PSF engineering, where commercially available DH phase masks (short-range: DH1-580, long-range: DH12R-670, both Double Helix Optics, Inc.)^39^ were aligned. In this work, fiducial beads emitting in a separate channel from the single molecules were used, enabling independent modulation of their emission with a long-range DH phase mask to accommodate longer acquisition times or correct for more severe drift.

### Sample preparation

For characterization of the soLLS, a coverslip spin-coated with donkey anti-rabbit secondary antibodies conjugated with dye CF568 (20098-1, Biotium) at a dilution of 1:10 in 1% (w/v) polyvinyl alcohol (PVA) prepared in nanopure water was used to visualize the cross-sectional profile of the soLLS and the Gaussian LS. The propagation profiles of the LSs were visualized using a microfluidic chip with a reflective mirror at a 45° angle filled with 0.2 mg/ml CF568 in nanopure water.

For quantification of the soLLS propagation in scattering environments, agarose powder (Sigma-Aldrich A9539) was dissolved in nanopore water to create a solution containing 1% (w/v) agarose gel. 10 µL of this gel was then mixed with 10 µL of 0.04 mg/ml CF568 in nanopure water and 10 µL of fluorescence beads (F8800, Invitrogen) at 1:10 dilution in nanopure water at 95°C and then dispensed into a microfluidic chip with a reflective mirror at a 45° angle right before imaging.

For cell experiments, human osteosarcoma cells (U-2 OS HTB-96, ATCC) were cultured and fixed inside microfluidic chips with a reflective mirror at a 39° angle. The microfluidic channels were cleaned with 70% ethanol (BP82031GAL, Fisher Scientific) for 10 min, followed by washing with phosphate-buffered saline (PBS) (SH3025601, Fisher Scientific) six times, and then coated with a 0.002% (w/v) solution of fibronectin (F0895, Sigma-Aldrich) in PBS for 2 h. U2OS cells were then seeded into the chips and incubated at 37°C and 5% carbon dioxide for 8 h before fixation. For fixation, cells were washed three times in PBS, fixed for 20 min in chilled 4% (w/v) formaldehyde (Electron Microscopy Science) dissolved in PBS, and then incubated for 10 min with 10 mM ammonium chloride (Sigma-Aldrich) in PBS.

For benchmarking the background reduction with soLLS, U2OS cells were labeled for lamin B1. After fixation, cells were permeabilized with three washes of 0.2% (v/v) Triton X-100 in PBS (5 min each), and subsequently blocked with 3% (w/v) bovine serum albumin (BSA) in PBS for 1 h. For both diffraction-limited and DNA-PAINT imaging of lamin B1, cells were incubated with rabbit anti-lamin B1 primary antibodies (ab16048, Abcam) at 1:1,000 dilution in 1% (w/v) BSA in PBS for 2 h, followed by three washes with 0.1% (v/v) Triton X-100 in PBS (3 min each). For diffraction-limited imaging, cells were incubated with donkey anti-rabbit secondary antibodies conjugated with CF568 (20098-1, Biotium) at 1:100 dilution in 1% (w/v) BSA in PBS for 1 h, then washed five times with 0.1% (v/v) Triton X-100 in PBS. For DNA-PAINT imaging, cells were incubated with donkey anti-rabbit secondary antibodies conjugated to oligonucleotides (Massive Photonics, order number AB2401012) at 1:100 dilution in antibody incubation buffer (Massive Photonics) for 1 h, followed by three washes with 1× washing buffer (Massive Photonics) in nanopure water. Complementary oligonucleotide-Cy3B dye conjugates (Massive Photonics, order number AB2401012) were diluted in imaging buffer (500 mM NaCl in PBS, pH 8) to a final concentration of 0.1 nM and 0.01 nM for 2D and 3D single-molecule imaging, respectively. For 3D imaging, the solutions were introduced into the chip at 50 mbar, corresponding to a flow rate of 7.4 µL/min, using a pressure-based flow control pump (LU-FEZ-0345, Fluigent, Inc.).

For 3D DNA-PAINT super-resolution imaging of TOMM20, LAP2, and lamin A/C, the cells were permeabilized in 0.1% (w/v) saponin (SAE0073, Sigma-Aldrich) in PBS for 10 min, and blocked with 10% (v/v) donkey serum and 0.05 mg/mL salmon sperm DNA in 0.1% (w/v) saponin in PBS for 1.5 h. For two-target TOMM20 and LAP2 imaging, the cells were labeled with rabbit anti-TOMM20 primary antibodies (ab186735, Abcam) and goat anti-thymopoietin (AF843, R&D Systems) primary antibodies at a dilution of 1:200 and 1:50, respectively, in 10% (v/v) donkey serum and 0.05 mg/mL salmon sperm ssDNA in 0.1% (w/v) saponin in PBS for 1 h. Cells were then washed three times with PBS (5 min each). After that, the cells were labeled with donkey anti-rabbit and donkey anti-goat oligonucleotide-conjugated secondary antibodies (Massive Photonics, order number AB2401012) at a dilution of 1:100 in antibody incubation buffer (Massive Photonics) for 1 h. For lamin A/C imaging, cells were labeled with mouse anti-lamin A/C primary antibodies (sc-376248, Santa Cruz Biotechnology) at a dilution of 1:100 in 10% (v/v) donkey serum and 0.05 mg/mL salmon sperm ssDNA in 0.1% (w/v) saponin in PBS for 1 h. After three washes with PBS, the cells were labeled with donkey anti-mouse oligonucleotide-conjugated secondary antibodies (Massive Photonics, order number AB2401012) at a dilution of 1:100 in antibody incubation buffer (Massive Photonics) for 1 h.

After labeling, the cells were washed three times with PBS and three times with 1× washing buffer (Massive Photonics) in nanopure water. Fiducial beads (F8807, Invitrogen) for drift correction were added to the chip at a dilution of 1:100,000 in PBS and incubated for 20 min before imaging. After fiducial beads were allowed to settle and attach, the cells were washed three times with 1× washing buffer.

During imaging, complimentary oligonucleotide-Cy3B dye conjugates (Massive Photonics, order number AB2401012) diluted in imaging buffer (500 mM NaCl in PBS, pH 8) were introduced into the chip at 100 mbar (corresponding to a flow rate of 14.9 µL/min) using the pump. For two-target imaging, the corresponding solutions were introduced at concentrations of 0.04 nM for TOMM20 and 0.08 nM for LAP2. For single-target imaging of TOMM20 or lamin A/C, corresponding solutions were introduced at a concentration of 0.04 nM.

### Acquisition settings

For all imaging, an EMCCD camera (iXon Ultra 897, Andor) was used for data acquisition. An EM gain set at 200 was used, corresponding to a calibrated EM gain of 182 for the EMCCD camera. The conversion gain of the EMCCD was experimentally determined to be 4.41 photoelectrons per A/D count. The EMCCD was operated at a shift speed of 3.3 µm/s with a standard vertical clock voltage amplitude. The readout rate was set to 17 MHz at 16-bit resolution using a preamplifier gain of 3. The calibrated pixel size of the camera was 159 nm/pixel vertically and 157 nm/pixel horizontally.

For characterization of the soLLS, image stacks of 200 frames of the cross-sectional views and 100 frames of the top and side views were acquired with the 560 nm laser at 0.2 W/cm^2^ and an exposure time of 100 ms. For the Gaussian LS, image stacks of 50 frames of the cross-sectional view and 100 frames of the side view were acquired with the 560 nm laser at 0.1 W/cm^2^ and an exposure time of 100 ms.

For propagation of the LSs in scattering environments, image stacks consisting of 500 frames were sequentially acquired from nine different FOVs for each condition (soLLS and Gaussian LS), using an exposure time of 100 ms. The 560 nm excitation laser was used at 7.2 W/cm^2^ for soLLS illumination and 13.6 W/cm^2^ for Gaussian LS illumination.

For diffraction-limited imaging of lamin B1, image stacks of 100 frames at an exposure time of 100 ms were acquired with the 560 nm laser at ∼1 W/cm^2^ for both soLLS and epi-illumination. For 2D single-molecule imaging of lamin B1, image stacks of 5000 frames at an exposure time of 200 ms were acquired with the 560 nm laser at ∼120 W/cm^2^ and ∼460 W/cm^2^ for soLLS and epi-illumination, respectively. For 3D single-molecule imaging of lamin B1, image stacks of 1000 frames at an exposure time of 200 ms were acquired with the 560 nm laser at ∼160 W/cm^2^ and ∼570 W/cm^2^ for soLLS and epi-illumination, respectively.

For 3D super-resolution imaging of TOMM20, LAP2, and lamin A/C, the 560 nm laser was used at 415 W/cm^2^ for soLLS illumination. For two-target imaging, image stacks of 60,000 frames were acquired for TOMM20 and 55,000 frames for LAP2 at an exposure time of 200 ms. For single-target imaging of TOMM20, image stacks of 150,000 frames were acquired at an exposure time of 200 ms. For single-target imaging of lamin A/C, image stacks of 50,000 frames were acquired at an exposure time of 200 ms.

### Data analysis

Diffraction-limited images were analyzed using ImageJ to extract intensity distributions and fitting results of the LS dimensions.

For comparing the soLLS and Gaussian LS in the scattering environments, image stacks of n = 9 different FOVs were acquired under soLLS and Gaussian LS illumination, and were cropped to the same size for post-processing. Image stacks were analyzed in ThunderSTORM^72^, an open-source ImageJ plugin, which localized the emitters in 2D, and provided the signal, background, and uncertainty of each localization. Data were processed with wavelet filtering (B-Spline order of 3 and scale of 2.0) and a weighted least-squares fitting. PSFs were detected using a maximum fitting method with a peak intensity threshold coefficient of 2. After filtering out low-confidence localizations (*vide infra*), the remaining localizations were binned into 5 rectangular regions of equal area for statistics comparison. The regions were labeled 1 to 5, with region 1 being closest to and region 5 being furthest away from beam focus.

For benchmarking the improvement in single-molecule imaging with soLLS, intensity line plots of the emitters were acquired from 2D single-molecule images of the same cell illuminated with soLLS or epi-illumination. The mean background reduction ratio between soLLS and epi-illumination were determined from intensity line scans over a 1.6 µm range where the background was measured, and the improvement in SBR were determined by averaging the mean ratio for n = 3 emitters for each cell. 3D single-molecule data for lamin B1 was analyzed with a modified version of Easy-DHPSF^73^, which localized the single-molecule data in 3D, and provided the signal, background, and 3D localization precision of each localization.

For 3D super-resolution imaging of TOMM20, LAP2, and lamin A/C, single-molecule data acquired with the short-range DH-PSF were localized using DECODE, an open-source deep learning-based software capable of localizing dense and overlapping emitters^31,42^. The DECODE model was trained using a z-scan of fluorescent beads imaged with the short-range DH-PSF and sparse 3D single-molecule data acquired from the same experimental setup, with setup-specific parameters: a dark level of 500 A/D counts, a conversion gain of 4.4 photoelectrons per A/D count, and an EM gain of 181.8. Training stopped after converging to a Jaccard index of 0.68 after 1000 epochs. The trained model was suitable for detecting signal photons in the range of 0–29,700 photons per localization and background levels of 0 to 240 photons per pixel. The trained model was then used for analyzing the various 3D single-molecule datasets throughout this work. The localizations were then drift-corrected using a custom MATLAB script with respect to the localizations of the fiducial beads within the FOV, which were acquired with the long-range DH-PSF and localized with a modified version of Easy-DHPSF.

### Filtering, rendering, and resolution analysis

Localizations were filtered to remove low-confidence localizations. For the soLLS versus Gaussian LS comparison in scattering environments, localizations with uncertainty larger than 100 nm or photon counts exceeding 1,000,000 photons per localization were discarded to remove outliers.

For 3D super-resolution imaging of TOMM20, LAP2, and lamin A/C, the localized data was filtered and rendered using the Vutara SRX (Bruker) software. For the two-target TOMM20 and LAP2 reconstruction, all TOMM20 localizations with localization precision greater than 42 nm in xy, greater than 50 nm in z, or having a signal photon count lower than 3,000 were removed. LAP2 localizations with a localization precision greater than 90 nm in xy, greater than 92 nm in z, or having a signal photon count lower than 500 were removed. For single-target multi-cell imaging of TOMM20, localizations with localization precision greater than 20 nm in xy or greater than 40 nm in z were removed. For single-target multi-cell imaging of lamin A/C, localizations with localization precision greater than 40 nm in xy or greater than 90 nm in z were removed. After filtering, all z localizations of the remaining localizations were scaled by a factor of 0.75 to account for index mismatch between the glass coverslip and the sample^31,63,74–76^.

The 3D super-resolution reconstructions were visualized by point splat rendering using a 25 nm particle size in Vutara SRX.

For resolution analysis, the FRC values were calculated in Vutara SRX using a super-resolution pixel size of 8 nm.

## Data availability

Source data for all graphs presented in this study are provided and will be made publicly available on GitHub upon completion of the peer review and publication process: [https://github.com/Gustavsson-Lab/soLLS-SourceData].

## Code availability

Analysis of 2D single-molecule data was performed using the open-source ImageJ plugin ThunderSTORM^72^ [https://github.com/zitmen/thunderstorm/releases/tag/v1.3/]. Data of short-range DH-PSF 3D super-resolution cell imaging with soLLS was analyzed using the open-source deep learning-based single-molecule localization software DECODE^42^ [https://github.com/TuragaLab/DECODE]. The modified version of Easy-DHPSF^73^ and the custom-written codes for drift correction are available on GitHub [https://github.com/Gustavsson-Lab/soLLS-SourceData].

## Acknowledgments

This work was supported by the National Institute of General Medical Sciences of the National Institutes of Health grant R35GM155365 and startup funds from the Cancer Prevention and Research Institute of Texas grant RR200025 to A.-K.G. The authors thank Yuya Nakatani for helpful discussions and assistance with data analysis. The microfluidic devices in this work were fabricated using resources and equipment available through the Shared Equipment Authority at Rice University.

## Author contributions

S.C. developed and constructed the imaging platform with assistance from all authors. S.C. and N.S. fabricated the microfluidic chips. S.C. prepared the samples and performed the experiments. S.C., N.S., G.G., and P.J. developed data analysis pipelines and S.C. analyzed the data. A.-K.G. conceived the idea and supervised the research. All authors contributed to writing of the manuscript.

