## Supplementary Fig. 1-9 and Supplementary Table 1 for "Single-objective lattice light sheet microscopy with microfluidics for single-molecule super-resolution imaging of mammalian cells"

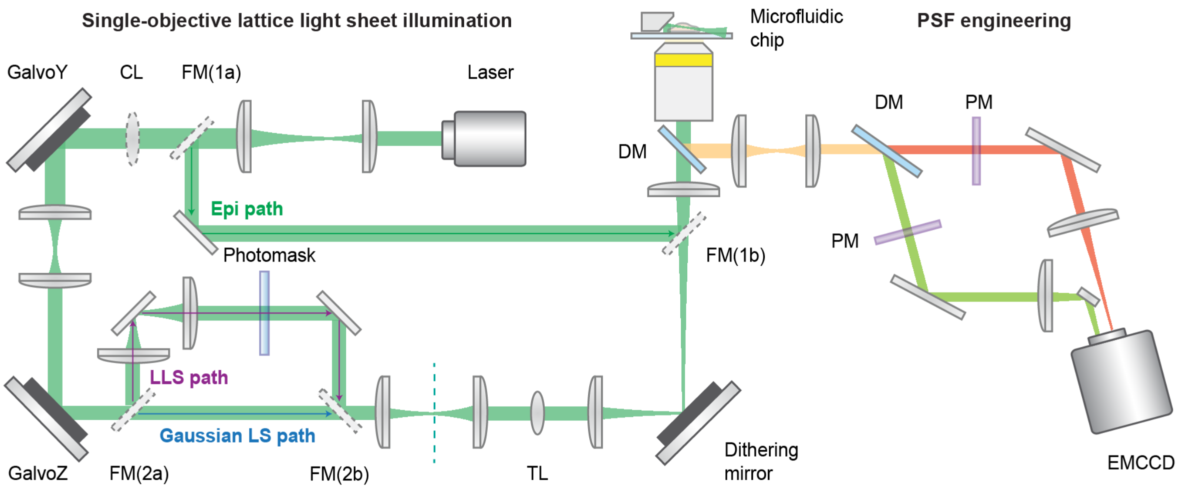


**Supplementary Fig. 1. Simplified schematic of the optical setup.** The single-objective lattice light sheet (soLLS) setup is coaligned with a single-objective Gaussian light sheet (LS) path and a widefield epi-illumination path. FM: flip mirrors that allow for switching between the three illumination modalities. CL: a cylindrical lens for generating the Gaussian LS. When FM(1a) and FM(1b) are flipped up, the setup works in epi-illumination mode. When FM(1a) and FM(1b) are flipped down and FM(2a) and FM(2b) are flipped up, the laser beam is shaped by the photomask to generate a LLS, and the setup works in soLLS illumination mode. When all the flip mirrors are flipped down, the laser beam is shaped by the CL, and the setup works in single-objective Gaussian LS illumination mode. GalvoZ, GalvoY, and TL: galvanometric mirrors and a tunable lens for LS steering. GalvoY is also used for scanning along the width of the LLS to homogenize its profile. These components are shared between the Gaussian LS path and the LLS path. Dithering mirror: mirror used to dither (adjust the angle of) the LS in the sample plane. DM: dichroic mirror. PM: phase masks in the emission paths for PSF engineering. The schematic is not to scale.

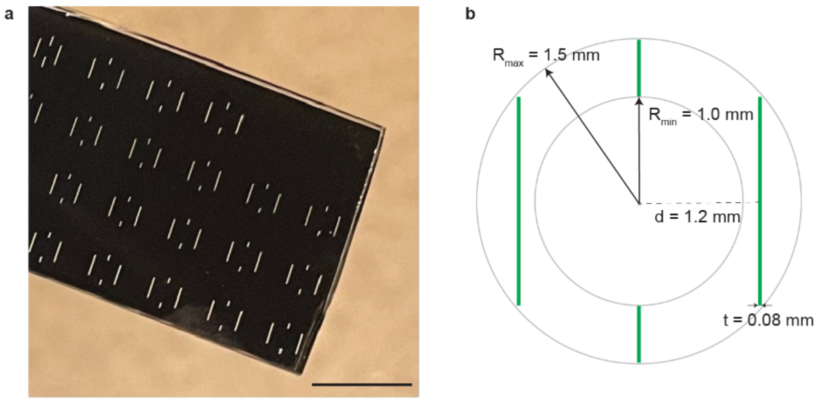


**Supplementary Fig. 2. Photograph and schematic of the photomask used in the soLLS platform. a,** A custom-designed photomask was implemented to generate the LLS. The print incorporates multiple mask pattern designs to facilitate flexible alignment adjustments. Scale bar: 1 cm. **b,** Dimensions of the slits pattern used in this work.

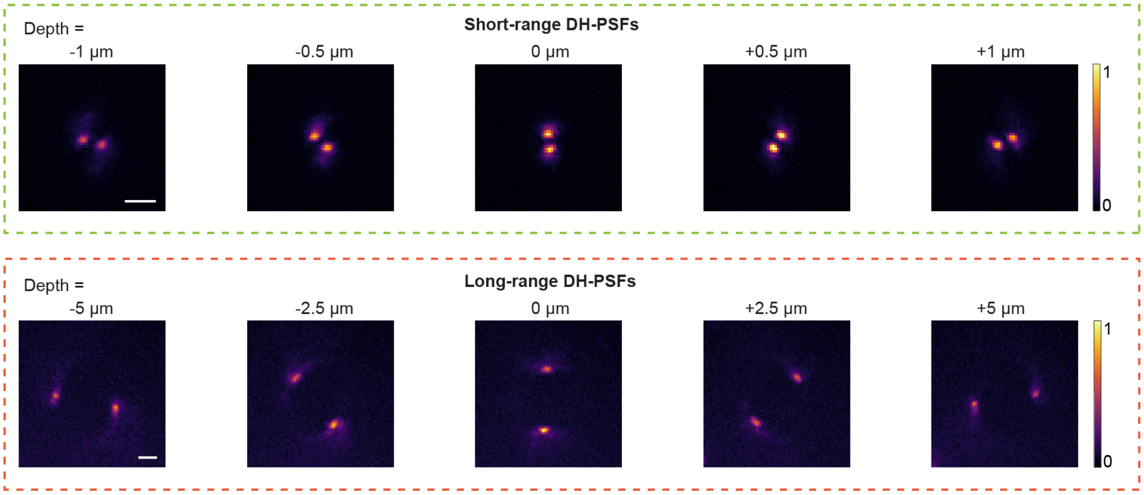


**Supplementary Fig. 3. Point spread function (PSF) engineering of the emission light yields 3D information.** In this work, short axial range double-helix (DH)-PSFs (top row), which have an experimental effective axial range of ~3 µm, was implemented for 3D localization of the single-molecule data. Long axial range DH-PSFs (bottom row), which have an experimental effective axial range of ~12 µm, were used for localizing fiducial beads for drift correction. Scale bars: 2 µm. The colorbars show normalized intensity.

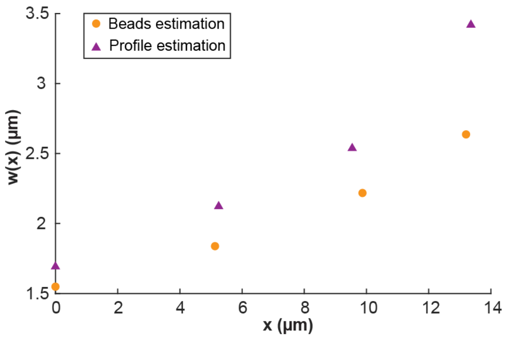


**Supplementary Fig. 4. Estimation of the thickness of the soLLS from intensity measurement of fluorescent beads.** Fluorescent beads were immobilized in 1% (w/v) agarose in a microfluidic chip with a mirror at 45°. The sample was scanned in 1D across the thin end (side view) of the soLLS. The thickness ($1/e^2$ radius) of the soLLS was estimated by fitting the intensity plot of the beads with a Gaussian function. The results show that the estimated beam thickness of the soLLS is smaller than the estimation from the projected profile captured using a CF568 solution in a microfluidic chip (Fig. 2c,f), as expected. Importantly, the trend of beam thickness during propagation remains consistent in both measurements.

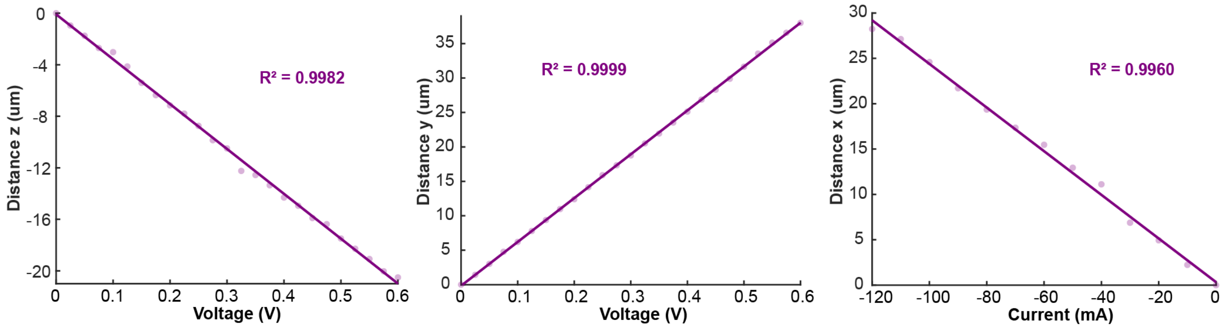


**Supplementary Fig. 5. Experimental calibration of the beam steering units in the soLLS setup.** The beam steering units featuring two galvanometric mirrors (GalvoZ and GalvoY) and a tunable lens (TL) to enable linear repositioning of the soLLS in three dimensions, where the LLS dimensions are decoupled from the steering. Experimentally-derived calibrations show that every 0.01 V applied to galvo Z/Y results in a translation in the z/y direction (up and down for sectioning and within the light sheet plane, respectively) of approximately 0.35/0.64 µm. The tunable lens shifts the light sheet focus 2.41 µm in the x direction (along the beam propagation) for every current step of 10 mA to enable sectioning of cells at various distances from the side wall.

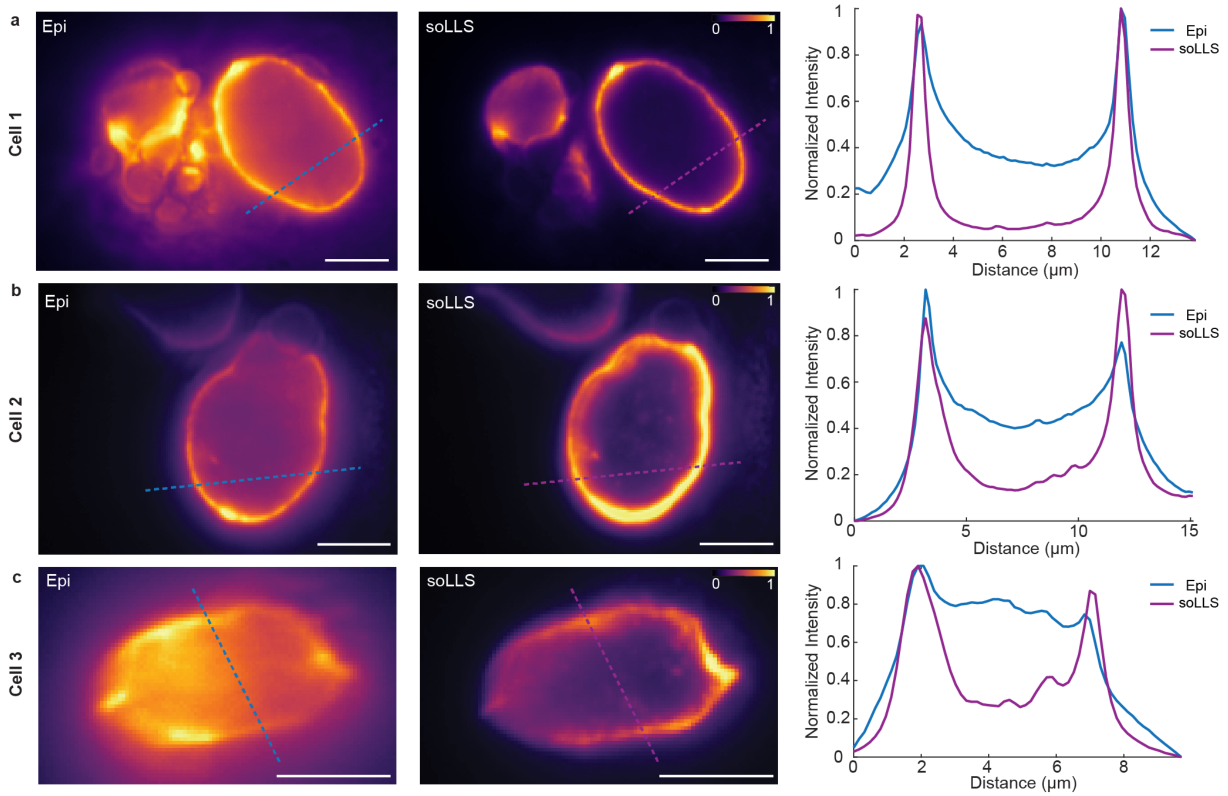


**Supplementary Fig. 6. Technical replicates demonstrating reproducibility of the diffraction-limited lamin B1 imaging shown in Figure 4a. a-c,** Diffraction-limited images of lamin B1 in three different U2OS cells acquired with widefield epi- (Epi) or soLLS illumination. Graphs show the comparison of normalized intensity distributions across line scans in the same cell under epi- or soLLS illumination, demonstrating reproducible signal-to-background ratio (SBR) improvement with soLLS illumination compared to widefield epi-illumination. SBR improvements were 7.6-fold, 3.0-fold, and 3.1-fold in **a**, **b**, and **c**, respectively. Scale bars: 5 μm. The colorbars show normalized intensity.

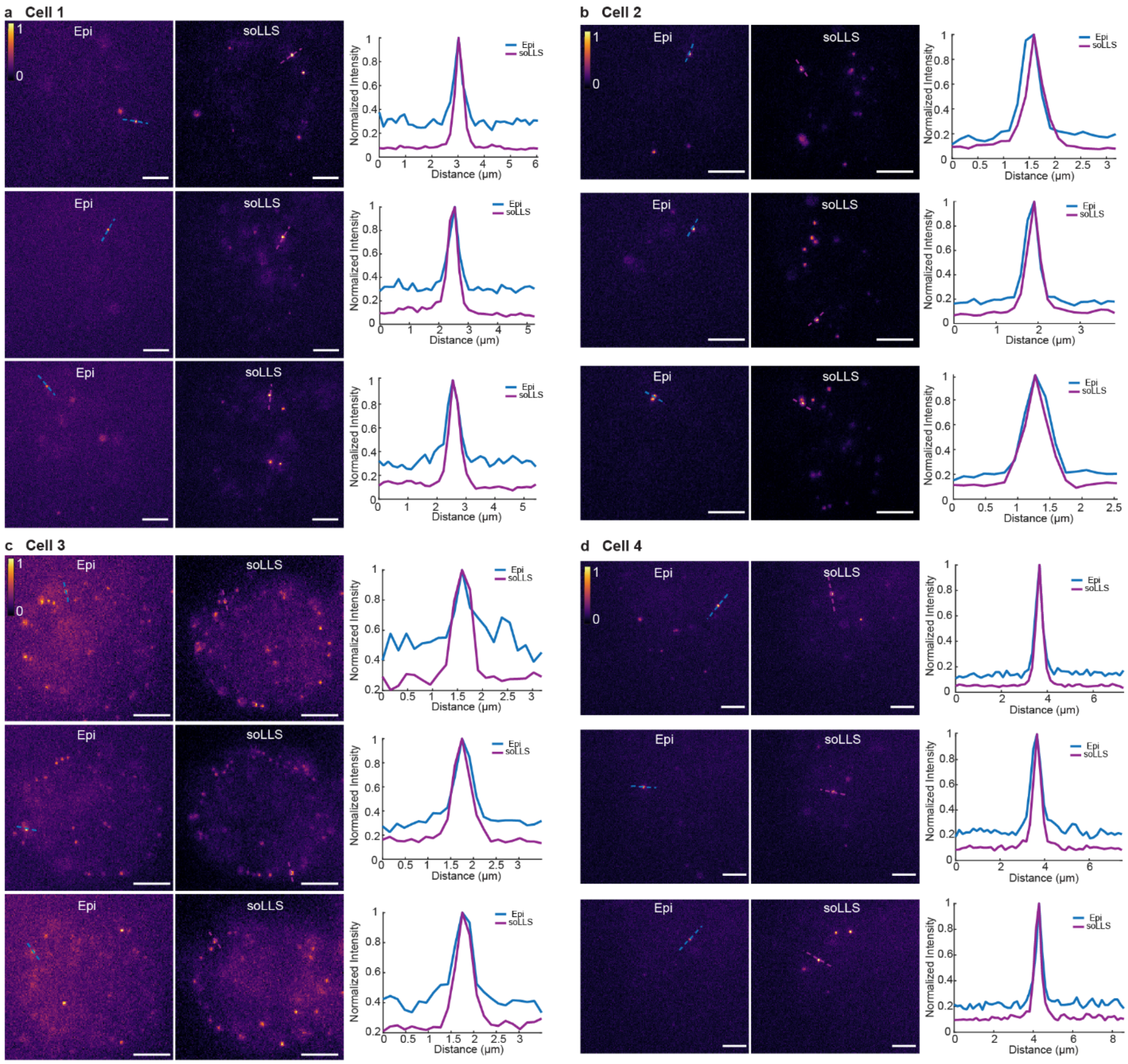


**Supplementary Fig. 7. Technical replicates demonstrating reproducibility of 2D single-molecule imaging of lamin B1 shown in Figure 4b. a-d,** 2D single-molecule images of DNA-PAINT labeled lamin B1 in four different U2OS cells acquired with widefield epi- (Epi) or soLLS illumination. Graphs show the comparison of normalized intensity distributions across line scans of the emitters under epi- or soLLS illumination. SBR improvements were 3.9±0.4-fold, 1.9±0.3-fold, 1.9±0.2-fold, and 2.5±0.5-fold in **a**, **b**, **c**, and **d**, respectively (mean ± standard deviation, n = 3 molecules per cell). Scale bars: 5 µm. The colorbars show normalized intensity.

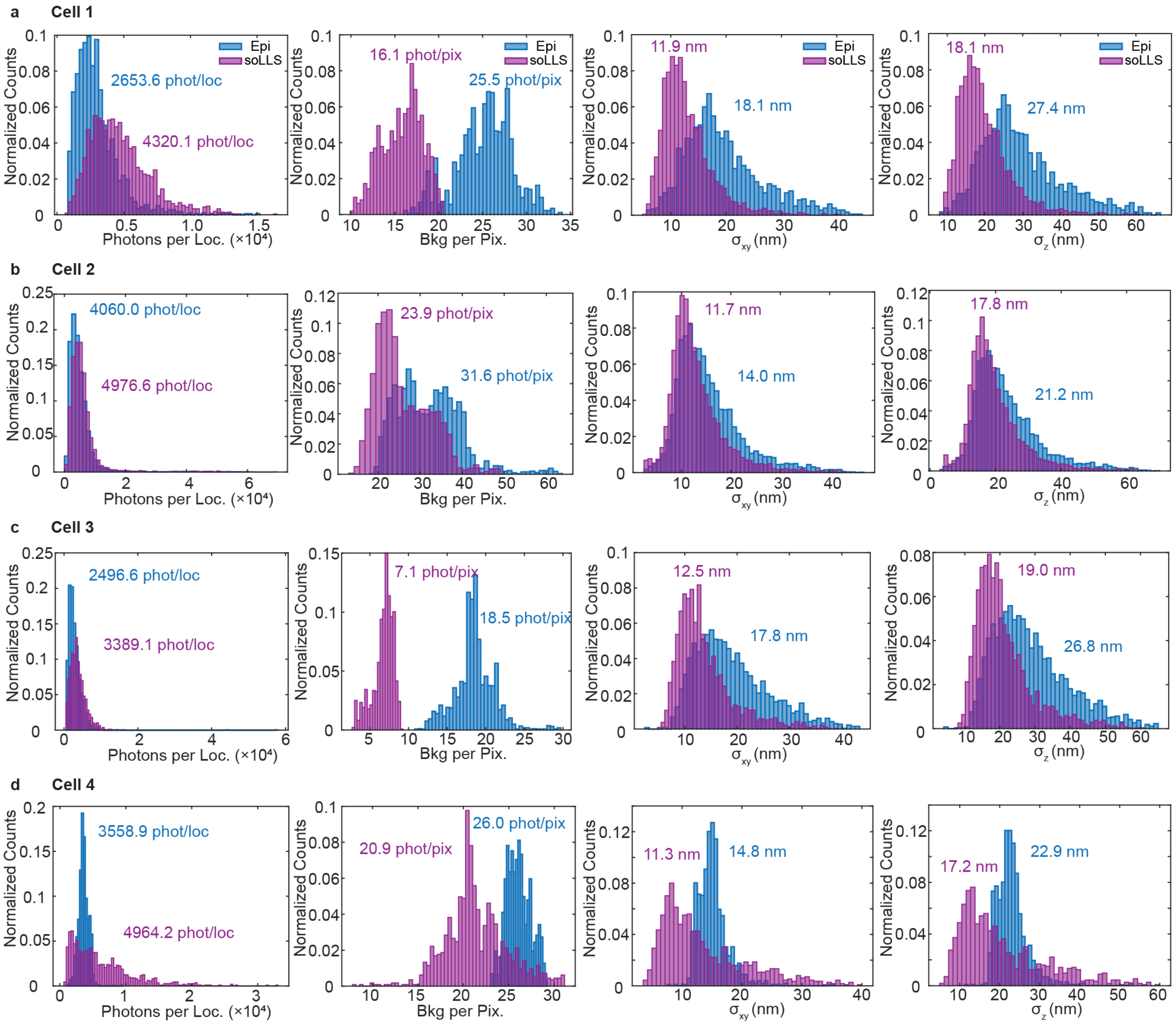


**Supplementary Fig. 8. Technical replicates demonstrating reproducibility of 3D single-molecule super-resolution imaging of lamin B1 shown in Figure 4c. a-d,** Histograms showing comparisons of signal photons per localization, background photons per pixel, lateral (xy) localization precision, and axial (z) localization precision of the localized emitters in the same cell under epi- (Epi) or soLLS illumination for four different cells. Median values of signal photons per localization, background photons per pixel, and localization precisions are indicated in each panel. Background reduction and improvements in localization precisions were demonstrated for each cell.

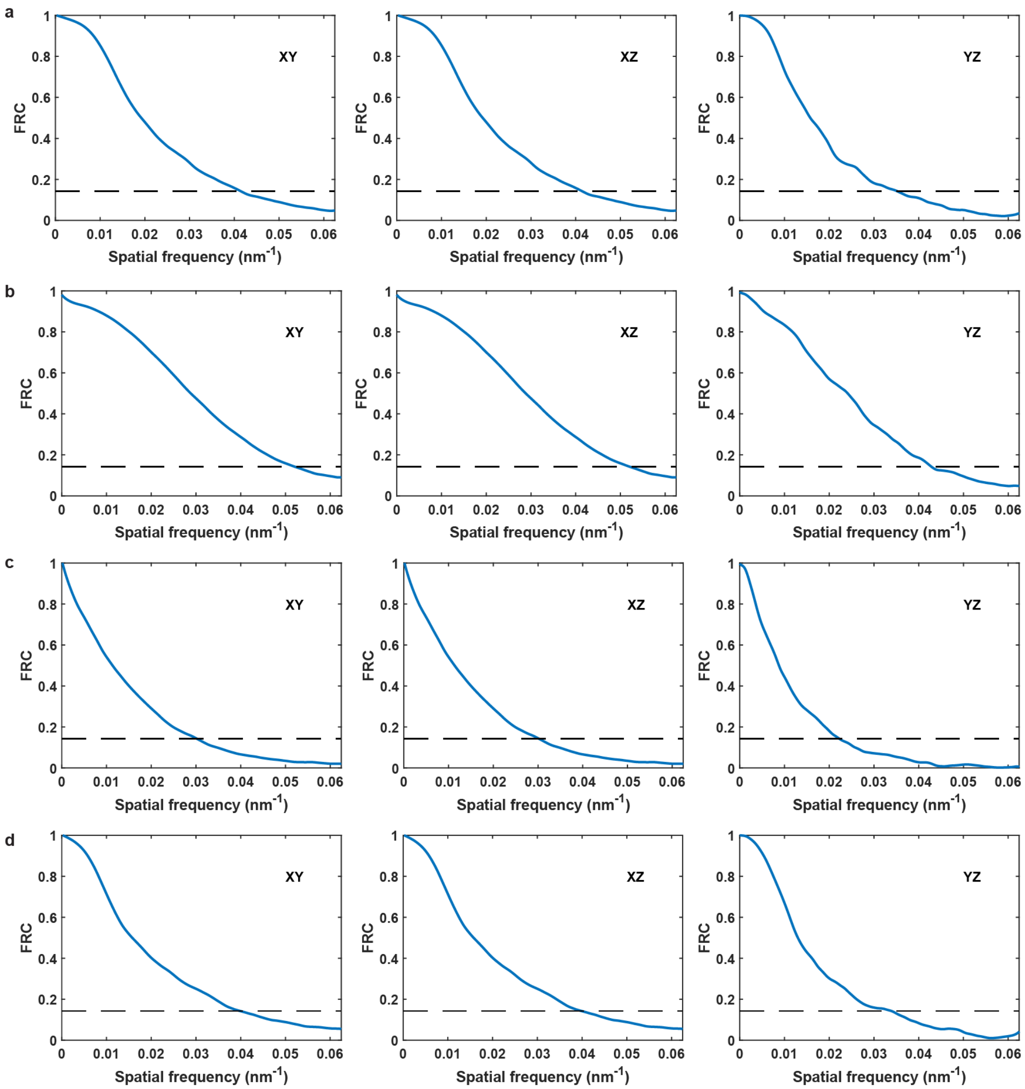


**Supplementary Fig. 9. Fourier ring correlation (FRC) analysis for the 3D super-resolution reconstructions shown in Fig. 5**. **a-b,** FRC analysis of the TOMM20 and LAP2 reconstructions shown in Fig. 5b-c, respectively. The resulting FRC resolutions in the xy/xz/yz planes were found to be **a,** 24.2/30.8/28.5 nm for TOMM20 and **b,** 19.2/23.4/23.4 nm for LAP2. **c-d,** FRC analysis of the lamin A/C and TOMM20 reconstructions shown in Fig. 5f-g, respectively. The resulting FRC resolutions in the xy/xz/yz planes were found to be **c,** 33.1/45.1/45.0 nm for lamin A/C and **d,** 25.1/33.6/29.5 nm for TOMM20. All FRC curves were calculated using a super-resolution pixel size of 8 nm in Vutara SRX.

**Supplementary Table 1. Absolute numbers of localizations in each region shown in Fig. 3.** The data is shown as mean ± standard deviation for n = 9 fields of view each for the two indicated illumination methods.

| | soLLS | Gaussian LS |
|---|---|---|
| Region 1 | 44488 ± 7820 | 49442 ± 13340 |
| Region 2 | 31426 ± 7444 | 12829 ± 8964 |
| Region 3 | 27087 ± 7861 | 2427 ± 2980 |
| Region 4 | 17989 ± 6802 | 746 ± 1341 |
| Region 5 | 7754 ± 4675 | 21 ± 22 |
